# Optimizing environmental conditions for PALM imaging of the nuclear pore complex labeled with the photoconvertible fluorescent protein mEos4b

**DOI:** 10.64898/2026.11.20.768092

**Authors:** Oarii Vahirua, Pascale Tacnet, Jérémy Néri, Anna Rosenkrancova, Jip Wulffelé, Oleksandr Glushonkov, Virgile Adam, Philippe Frachet, Bernhard Brutscher, Dominique Bourgeois

**Author notes:** Corresponding Author Dominique Bourgeois − Univ. Grenoble Alpes, CEA, CNRS, Institut de Biologie Structurale (IBS), 38000 Grenoble, France.

## Abstract

Green-to-red photoconvertible fluorescent proteins (PCFPs) are smart fluorophores used in several nanoscopy approaches based on single-molecule localization microscopy (SMLM). To optimize the achievable spatial resolution and facilitate molecular counting in these approaches, PCFPs should deliver a high effective labeling efficiency (ELE). Here, to evaluate the ELE provided by mEos4b, a performant PCFP, we designed a constitutive Nup96-mEos4b protein fusion in the nuclear pore complex (NPC) of U2OS cells, and performed PhotoActivated Localization Microscopy (PALM) imaging of NPCs in fixed cells under different environmental conditions. We find that removal of oxygen improves the ELE by more than 20%. Overall, using HEPES as buffer in deuterated and deoxygenated water delivers the highest quality PALM images of NPCs labeled with mEos4b. These conditions should be beneficial for quantitative PALM or Minimal Fluorescence Photon Fluxes Microscopy (Minflux) experiments in fixed cells. We propose a photophysical scheme that accounts for the observed improvements in the absence of oxygen.

---

Photoconvertible fluorescent proteins (PCFPs) have remained attractive fluorophores for advanced fluorescence microscopy approaches over the past decade.^1,2^ Like other fluorescent proteins (FPs), PCFPs allow genetically-encoded labelling, resulting in defined stoichiometry between the protein of interest and the fluorescent label, and high target specificity without the need for exogenous labeling reagents. Members of the EosFP family^3^ such as mEos3.2, mEos4b, pcStar or mEosEM stand out due to their high photon budget, good monomeric quality, and, for mEos4b and mEosEM, strong fixation resistance. Recently, fast maturing variants of mEos4b have been introduced,^4^ further expanding the applicability of these probes. These properties make EosFP-derived PCFPs well-suited for quantitative nanoscopy experiments at the single-molecule level.^5^

Green-to-red PCFPs are initially in a green-emitting form and can be converted to an orange-red emitting form upon absorption of a violet photon (∼405 nm). This process occurs with a low photoconversion quantum yield, on the order of 1e^-4^.^6^ A conserved feature of these proteins is a histidine at the first position of the chromophore triad (His62 in mEos4b). Photoconversion involves backbone cleavage between this histidine and the preceding residue Phe61, thereby extending the chromophore’s electron conjugation to the imidazole ring of His62. The underlying photoconversion mechanism, however, remains incompletely understood.^3^ Beyond photoconversion, EosFP-derived PCFPs exhibit complex photophysical behavior, including multiple forms of fluorescence blinking and irreversible bleaching.^3,7^ More recently, ground-state heterogeneities in both the green^8^ and the red forms^9^ have been reported. Overall, these PCFPs display highly intricate photophysical properties.

In PhotoActivated Localization Microscopy (PALM), these photophysical properties directly influence one essential parameter: the effective labeling efficiency (ELE).^10^ The ELE is the fraction of PCFPs that is successfully detected during imaging and is dependent on a combination of factors, including efficiency of fluorescent-protein maturation, photo-conversion kinetics, and the photophysical properties of both the green- and red-emitting forms. Achieving a high ELE is essential for obtaining high-resolution images and for accurate molecular counting.

Because the photophysical properties of PCFPs can be strongly influenced by their local physicochemical environment,^11^ characterizing the ELE *in cellulo* is essential for assessing their performance under biologically relevant conditions. The nuclear pore complex (NPC) has emerged as a widely used metrology standard for evaluating labeling strategies and image-acquisition protocols.^10,12–14^ In their pioneering study, Thevatasan et al^12^ demonstrated high-quality PALM imaging using the PCFP mMaple^15^ fused to the nucleoporin Nup96, with TRIS buffer at pH 8 and heavy water (D_2_O).^16^

Each NPC contains 32 copies of Nup96, organized into two rings, each comprising eight symmetrically arranged Nup96 pairs. Therefore, counting the number of visible corners for each detected NPC in 2D images provides a reliable estimate of the ELE.^12^

Here we generated a Nup96-mEos4b stable human bone osteosarcoma (U2OS) cell line, using a CrispR-Cas9 genome-editing strategy similar to a previously described approach^17^ (Fig. S1).

We first compared the performance of mEos4b to that of mMaple, originally employed by Thevatasan et al^12^ for PALM imaging of NPCs in fixed cells, and using the same environmental conditions, ie TRIS buffer at pH 8 and heavy water (D_2_O) (Fig. 1, Fig. S2). We evaluated four complementary metrics: localization precision (LP), effective labeling efficiency (ELE), number of photons per localization (Nphot) and image resolution quantified using Fourier ring correlation (FRC).^18^ Although these metrics are interrelated, each captures a distinct aspect of imaging performance (Supporting Text).

**Figure 1:**
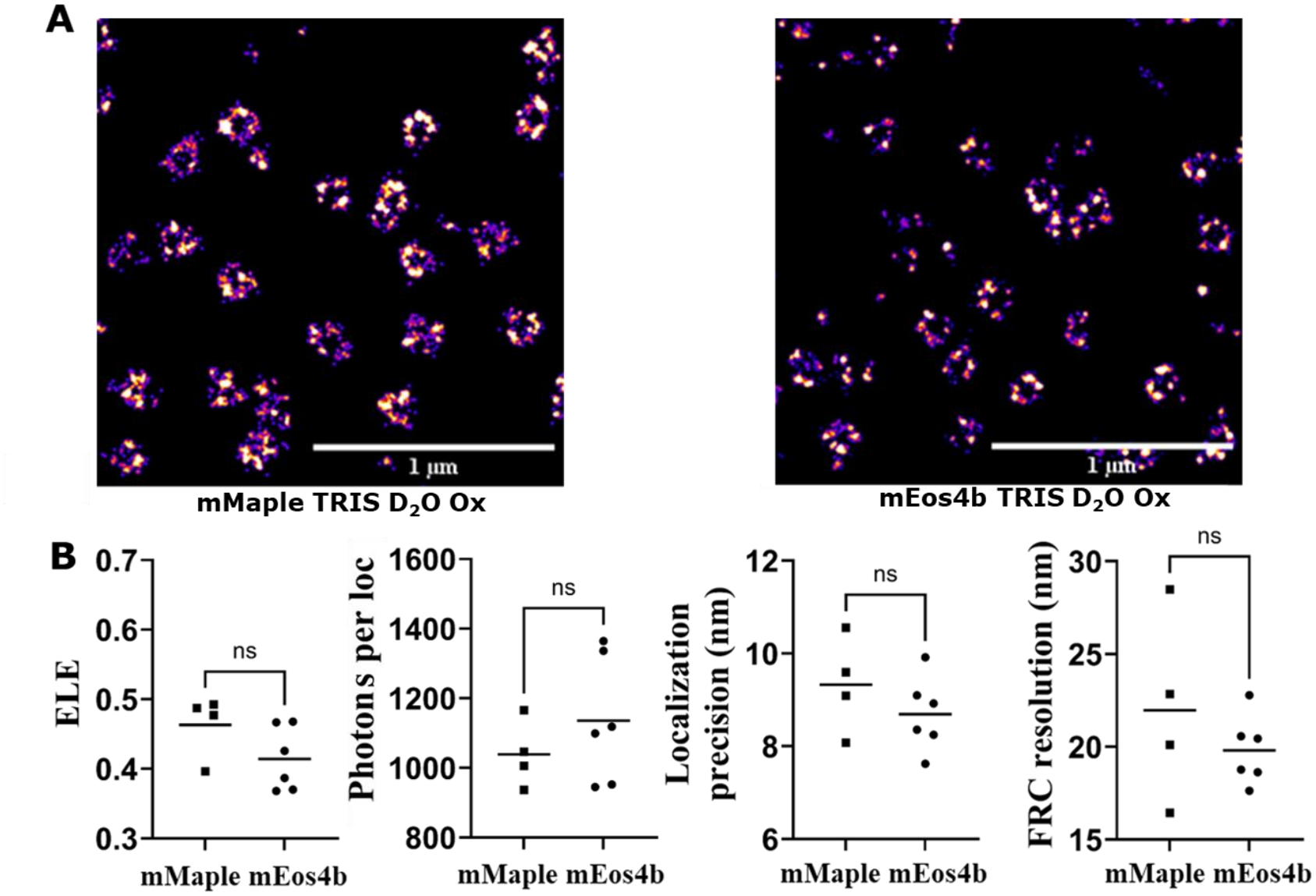
Comparison of the performance of mMaple and mEos4b fused to Nup96 in PALM under reference environmental conditions. **(A)** Representative PALM reconstructions of NPCs in fixed U2OS cells expressing Nup96-mMaple (left) or Nup96-mEos4b (right). Data acquired under identical conditions. Scale bars, 1 µm. **(B)** Comparisons of mean effective labeling efficiency (ELE), localization precision, number of photons per localization and resolution based on Fourier ring correlation (FRC) between mMaple and mEos4b. For each independent experiment, at least three spatially distinct fields of view (FOVs) were acquired and analyzed. Each parameter was determined independently for each FOV, and the resulting values were averaged to obtain a single coverslip-level measurement. Each symbol represents one independent experiment (one coverslip; mMaple, n = 4; mEos4b, n = 6), and horizontal lines indicate group means. Statistical comparisons were performed using two-tailed, unpaired Mann–Whitney U tests. ns, not significant (p > 0.05).

Based on our data processing protocol (Supporting Materials & Methods), no significant differences were observed between the two PCFPs in our NPC images. Both FPs yielded median LPs below 10 nm, ELEs between 40 and 50% and FRC resolutions of approximately 20 nm. Thus, the reported ∼30% superior brightness and higher photon budget reported for mEos variants compared with mMaple and Dendra variants in their red forms *in vitro*^19^ were not apparent in our data. This discrepancy may reflect the masking of in vitro differences by cellular variability or changes in photophysical properties induced by the cellular environment.

Next, we changed the environmental conditions for NPC image acquisition to investigate their effects on the quality of our data sets. We first confirmed that heavy water has a beneficial effect (Fig. S3),^12,16^ in agreement with previous work reporting a gain in PCFP photostability and brightness.^16^ Then, we modified the buffer and tested HEPES (pH 7.5), a more cell-friendly buffer as compared to TRIS (pH 8.0).^21,22^ The use of HEPES was also motivated by the notion that, in view of the red-form heterogeneity of mEos4b reported recently,^9^ HEPES buffer favors population of the brighter red form as compared to TRIS. However, no effect on Nphot was observed, while a significant ∼20% improvement was obtained for the ELE. Using a universal buffer^23^ to decouple the effects of buffer (TRIS vs HEPES) and pH (8.0 vs 7.5) (Fig. S4) suggested that the lower pH of HEPES could be partly responsible for the observed higher ELE, although this difference is statistically not significant, most likely because of cellular variability. Overall, these results suggest that buffer composition, in addition to pH, may influence data quality. Among the conditions tested, HEPES at pH 7.5 appears to be the preferred choice.

In contrast to tuning buffer or pH, removal of oxygen using a glucose oxidase / catalase / glucose (GLOX) system provided a clear improvement in both ELE, and Nphot, translating into high-quality rendered images with many well-defined 8-corners NPC rings (Fig. 2). The FRC resolution was not improved, in line with simulations using SMIS (Fig. S5, Supporting Text).^24^ More surprisingly, unlike the photon yield, localization precision did not improve. This is attributed to the consistently higher fluorescence background observed in the presence of GLOX (Fig. S6), degrading the LP in line with the Thompson formula.^25^ The newly introduced oxygen scavenger Na_2_SO_3_^26^ was also tested and provided similar improvement (Fig. S7). Molecular oxygen is known to diffuse into the chromophore pocket of practically all FPs and to efficiently quench the triplet state (T_1_) through the formation of reactive oxygen species (ROS). ROS such as singlet oxygen (^1^O_2_) may then react with the FP scaffold, resulting in photobleaching. Thus, oxygen removal is expected to photostabilize mEos4b, consistent with the observed improvements in image quality.

**Figure 2:**
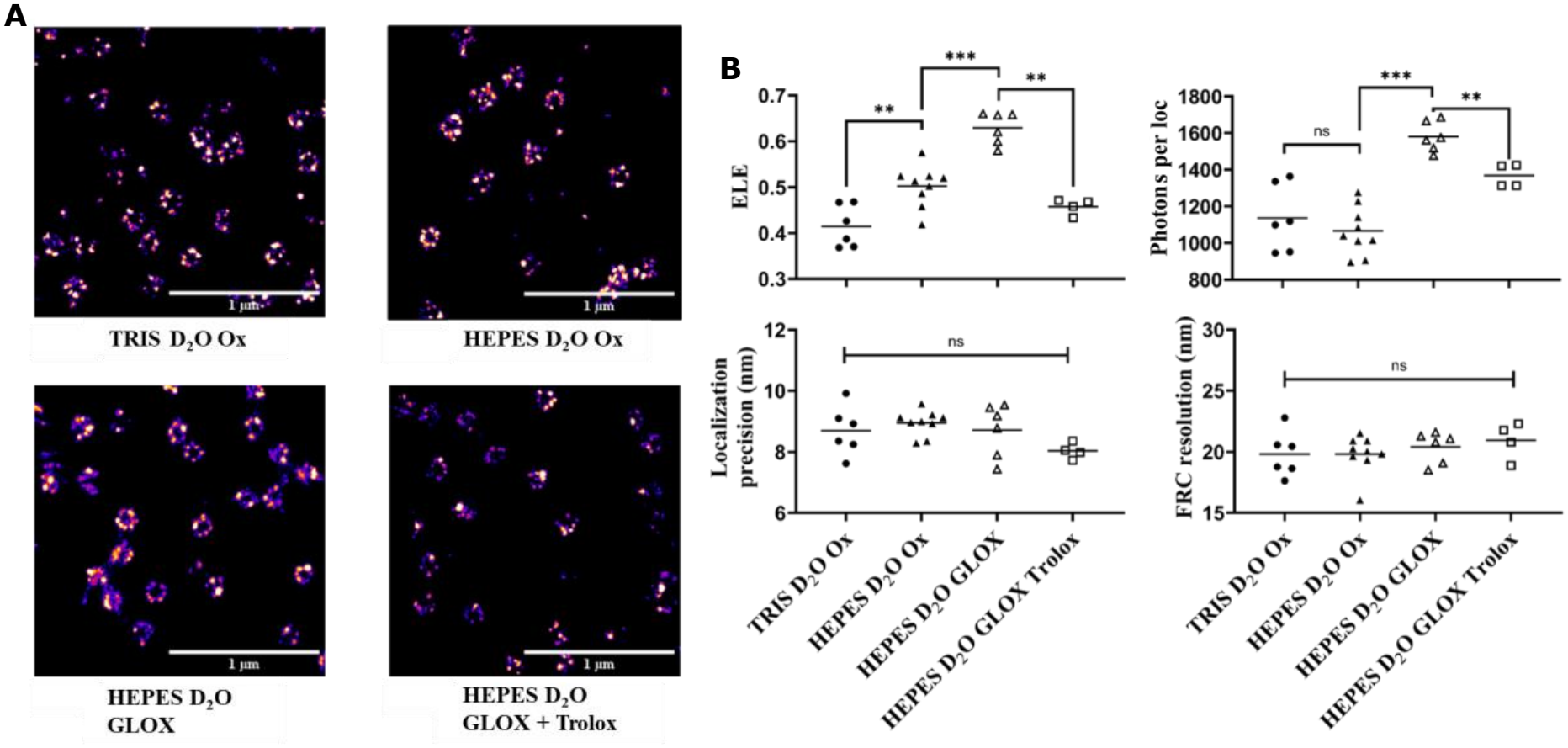
Optimization of environmental conditions for PALM imaging of Nup96-mEos4b in fixed cells. **(A)** Representative PALM reconstructions of NPCs in fixed U2OS Nup96-mEos4b cells acquired under four environmental conditions: Tris buffer (pH 8.0) in D_2_O under standard oxygenated conditions; HEPES buffer (pH 7.5) in D_2_O under standard oxygenated conditions; HEPES buffer (pH 7.5) in D_2_O supplemented with a glucose oxidase/catalase (GLOX) oxygen-scavenging system; and the same HEPES/GLOX buffer supplemented with 1 mM Trolox. Scale bars, 1 µm. **(B)** Comparisons of effective labeling efficiency (ELE), median localization precision, median number of photons per localization and resolution based on Fourier ring correlation (FRC) across the four conditions. For each independent experiment, at least three spatially distinct fields of view (FOVs) were acquired and analyzed. Each parameter was determined independently for each FOV, and the resulting values were averaged to obtain a single coverslip-level measurement. Each symbol represents one independent experiment (one coverslip), and horizontal lines indicate group means. Statistical comparisons were performed using two-tailed, unpaired Mann–Whitney U tests. Statistical significance is indicated as * (p < 0.05), ** (p < 0.01) and *** (p < 0.001). ns, not significant (p > 0.05).

To address whether the increase of ELE observed under anaerobic conditions arises from enhanced photostability of the red form, of the green form, or from a combination of both, we performed solution NMR spectroscopy combined with in situ sample illumination.^27^ We quantified the formation of the red form of mEos4b upon 405 nm irradiation in the presence and absence of O_2_ (Fig. 3). Under the NMR experimental conditions used (see Supporting Materials and Methods), the total amount of red mEos4b increased in the absence of O_2_ by a proportion comparable to the increase in ELE observed at the single-molecule level. These results support the interpretation that the increase in ELE under anaerobic conditions arises predominantly from beneficial changes in the photophysics of the green form.

**Figure 3:**
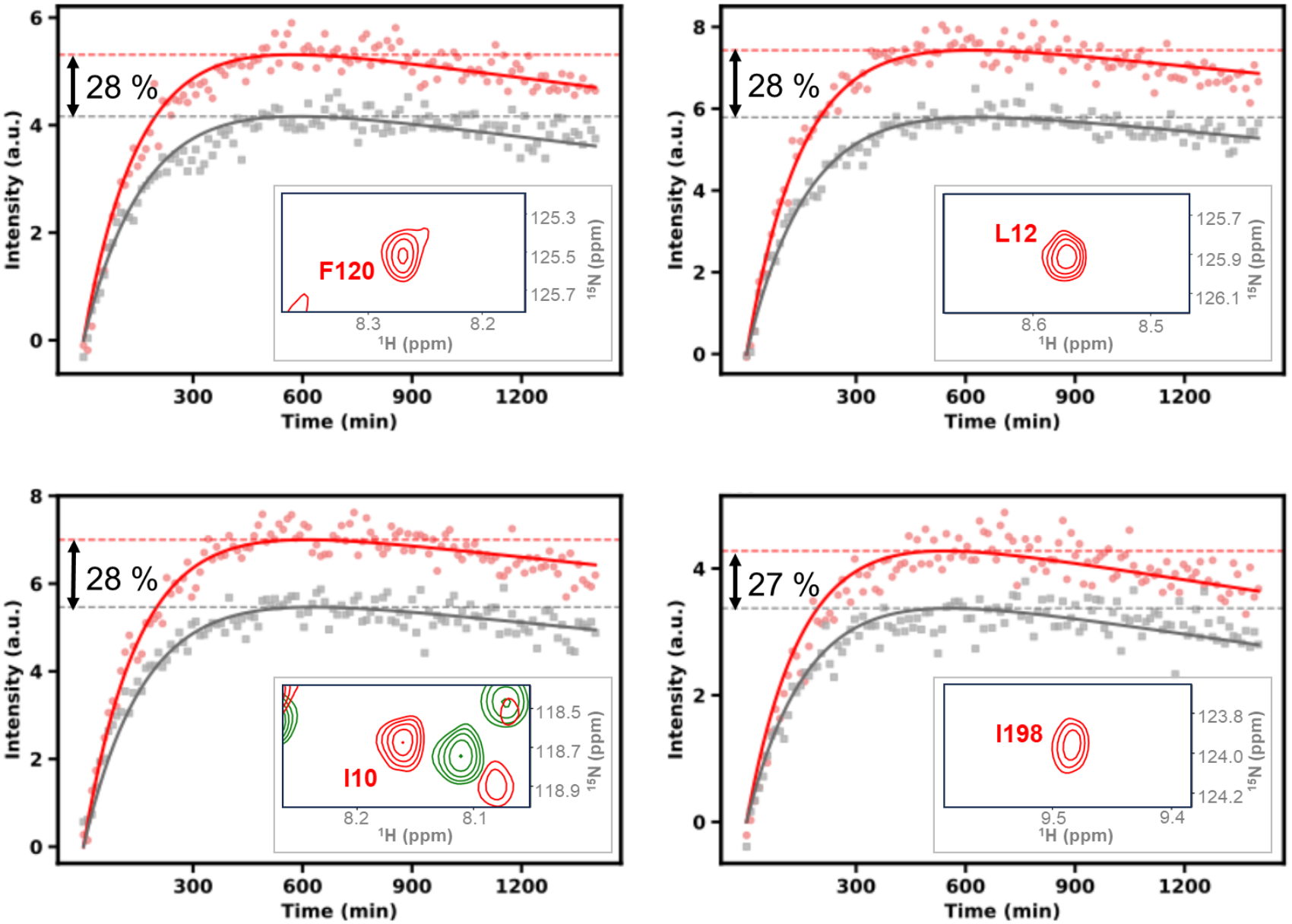
mEos4b photoconversion kinetics monitored by NMR spectroscopy in the presence or absence of oxygen. NMR signal intensities from photoconverted red mEos4b were extracted from four representative residues (F120, L12, I10 and I198) exhibiting well resolved NMR peaks (insets) and plotted as a function of illumination time. The red-state population build-up curves obtained under oxygen-depleted (red dots) and oxygen-rich (gray squares) conditions are shown. Solid lines (red and grey) represent best fits to a function combining mono-exponential growth with linear signal decay and are intended solely to guide the eye. The maximum red-state population, reporting on the photoconversion yield, increased by 28 ± 4% upon removal of molecular oxygen from the sample solution.

We also attempted to collect PALM data in the absence of oxygen on live cells. We systematically observed a drastic reduction in ELE, while the photon budget per localization and the localization precision remained superior to those measured in oxygenated conditions (Fig. S8). Clearly, living cells rapidly suffer from anoxia, likely giving rise to NPC disruption.^28^ These data highlight that environmental optimization needs to consider cellular integrity in addition to photophysical performance.

We then recorded PALM data on fixed cells using TROLOX in addition to GLOX. TROLOX has been shown to exhibit both oxidizing and reducing properties,^29^ efficiently acting on radical states forming downstream T_1_ in some organic dyes through the so-called ROXS effect.^30^ As short-lived radicals may also form in FPs and electrons or holes may diffuse out of the β-barrel, a beneficial effect of TROLOX could also be expected with mEos4b in PALM. However, the opposite was observed, as both the ELE and localization precision degraded (Fig. 2). Nevertheless, these data suggest that TROLOX, which is unlikely to diffuse within the FP scaffold, exerts a significant indirect effect on mEos4b photophysics, possibly through radical stabilization.

Finally, we investigated whether the improvements in PALM imaging of NPCs in fixed mammalian cells resulting from oxygen removal could be confirmed in a non-biological context using *in vitro* single-molecule measurements. For this purpose, we immobilized mEos4b using a Spy-catch Spy-tag matrix, as described recently.^19^ We then analyzed photophysical red-form properties of mEos4b in the presence or absence of O_2_ and TROLOX. Although the ELE could not be measured in the absence of labeling to a template of well-defined stoichiometry, reliable single-molecule fluorescence traces could be extracted. This was not possible *in cellulo*, where the local density of activated mEos4b around individual NPCs is typically too high. Analysis of the full fluorescence traces provided access to several parameters, including the total photon budget, on- and off-times, and number of blinking events (excursion to transient dark states), in addition to the number of photons detected per localization. The results, presented in Fig. S9, confirm that the photostability of red mEos4b is substantially enhanced in the absence of molecular oxygen. Importantly, the number of photons emitted per localization before photobleaching (see Materials & Methods) remained higher under anaerobic as compared to oxygenated conditions. This finding, which is consistent with the increased photon yield observed *in cellulo* in the absence of O_2_ (Fig. 2), strongly suggests that oxygen depletion enhances not only photostability but also apparent brightness. This observation is somewhat unexpected in view of the anticipated increase in T_1_ lifetime^31^ and therefore stronger T_1_ saturation in the absence of O_2_. Finally, we observed a modest but significant reduction in on-times and an increase in off-times upon oxygen removal. These changes are mainly attributed to an increase in the number of long-lived blinking events (associated with cis-trans isomerization of the chromophore) relative to short-lived events (associated with radical formation), but may also result from a small lengthening of those latter events, as anionic radicals could also be oxidized by molecular oxygen.

Overall, we propose the photophysical model of Fig. 4 to explain the beneficial effect of O_2_ removal on PALM imaging. SMIS simulations^24^ of NPC PALM data based on this model reproduced the experimental findings (Fig. S10). According to the model, oxygen removal enhances the ELE through improved photostability of both the green- and red-emitting forms of mEos4b.

**Figure 4:**
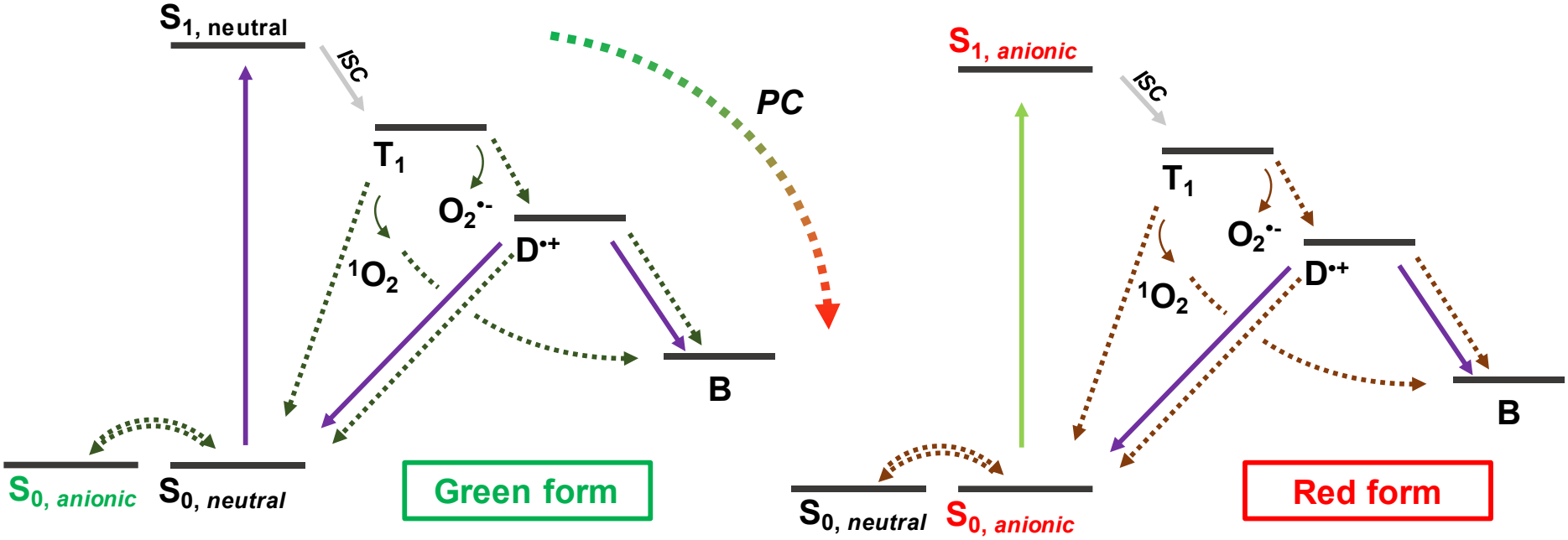
Proposed photophysical scheme accounting for increased brightness and photostability of mEos4b upon oxygen removal in both its green and red states. S: Singlet state, T1: Triplet state, B: Bleached, D^•+^: Cationic radical, PC: photoconversion. Plain colored lines: light-induced transitions, dotted lines: thermally-induced transitions. For a complete photophysical model, see Fig. S10.

The contribution of O_2_ removal to green-form photostabilization is supported by the NMR measurements. These experiments quantify photoconversion kinetics under weak 405 nm illumination alone, without additional 561 nm excitation, thereby minimizing the contribution of red-form photophysics to the observed photoconversion efficiencies. NMR provides a quantitative readout of molecular-state populations, as the integrated volumes of NMR signals are proportional to the populations of the corresponding states under appropriate acquisition conditions. We therefore fitted the measured photo-conversion kinetics to a simplified model derived from Fig. 4 (Fig. S11), that also accounts for the experimentally observed green- and red-form heterogeneities.^8,9^ The resulting fits (Fig. 3 and Fig. S11) are consistent with an increased population of the red-emitting form under anaerobic conditions (∼28%), matching well the increase in ELE measured in PALM (∼23%). Within the framework of the model, the improved photoconversion efficiency can be attributed to an approximately 40% reduction in the photobleaching quantum yield of the green-emitting form.

Red-form photostabilization, on the other hand, is directly supported by the single-molecule fluorescence measurements, as evidenced by the increased number of photons per localization in NPC PALM experiments and the higher total photon budget per mEos4b molecule *in vitro*.

In both green and red forms, O_2_ is responsible for ROS-mediated photobleaching through triplet-triplet annihilation with T_1_. Released singlet oxygen (through the so-called type II photochemical pathway), due to its short mean free path, may efficiently oxidize amino acids of the PCFP scaffold. In particular, sulfoxidation of methionine and cysteine residues near the chromophore may drastically rise its pKa, locking the chromophore in a nonfluorescent protonated state.^32^

In the green form, because photobleaching and photoconversion are competing processes, a reduction in photobleaching translates into an increase in the overall photoconversion yield.

The NMR data also suggest that the intrinsic rate of photoconversion does not substantially change upon O_2_ removal. This points toward the hypothesis that T_1_, the lifetime of which is expected to increase in anaerobic conditions, may not be directly involved in “standard” photoconversion,^33^ in contrast with “primed” photoconversion occurring in PCFPs such as Dendra2 or mMaple3.^34^

The observation that the number of photons recorded per localization increases moderately before irreversible photobleaching occurs, substantiated by the *in vitro* data, is unexpected. Indeed, in the absence of O_2_, triplet state saturation increases due to reduced quenching. Given the low intersystem crossing (ISC) quantum yields of 0.1 to 1% and millisecond-range T_1_ lifetimes reported for FPs^31^, triplet state saturation can substantially limit apparent brightness under single-molecule imaging conditions, which typically require high excitation laser power densities. This limitation would therefore be expected to become even more pronounced under anaerobic conditions. The paradoxical increase in apparent brightness observed upon oxygen removal thus suggests the involvement of an additional, downstream dark state with a short life time relative to the camera frame time, whose formation is suppressed under anaerobic conditions. Such a dark state could be a cationic radical resulting from O_2_-mediated T_1_ oxidation (ie type I photochemical pathway), as described previously in the case of organic dyes.^30^ Further studies will be required to establish whether such a radical state also occurs in PCFPs.

Another conclusion of our study is that optimization of environmental conditions for quantitative PALM should preferably be performed *in cellulo* to maximally approach real biological conditions. Nevertheless, cellular variability can mask the statistical significance of certain differences between photophysical behaviors of PCFPs. Furthermore, multiple independent measurements are required to minimize the risk of drawing false conclusions, highlighting the need for high-content-screening (HCS) approaches^19^ to compare the photophysical performance of multiple PCFPs under fixed environmental conditions *in vitro*. An additional essential aspect of performing optimization in cells is that physicochemical parameters that favor superior photophysical performance may also be assessed in terms of their possible deleterious effects on cellular integrity, as observed here with live cells not withstanding oxygen removal. Moreover, subtle findings may emerge from *in cellulo* data, such as the fact that O_2_ removal, while increasing PCFP’s photon yields, increases background fluorescence, reducing the expected gain in localization precision. This finding may not be surprising as the photostability and/or brightness of fluorescent molecules contributing to the background signal (including out-of-focus PCFPs) may also increase in oxygen-free conditions. In general, a limitation of *in vitro* studies is that specific fusion constructs or the local environment in cells may modify the photophysical behavior of FPs despite the caging effect of the β-barrel. This aspect is exemplified here with the observed detrimental effect of TROLOX when added to GLOX, a molecule that is unlikely to penetrate the FP protein scaffold but still significantly alters photophysical performance.

The ELE is a critical parameter for maximizing the effective resolution of reconstructed super-resolution images and ensuring accurate molecular counting. Measuring ELE *in vitro*, however, requires sophisticated experimental strategies.^5^ In contrast, the NPC provides a convenient reference for assessing the ELE *in cellulo*, as exemplified by numerous studies exploiting this well behaved molecular template. Nevertheless, absolute ELE values should be considered with caution, as they depend on imaging parameters and the selected data-processing pipeline (Supporting Text). Accordingly, relative ELE values obtained from comparable experiments are likely to provide more robust and reliable measures.

In the present study, the ELE reached a maximum of approximately 60% under our optimized conditions. This implies that about 40% of the molecules were either excluded during data processing because of insufficient localization precision or remained undetected, due to lack of maturation, photobleaching in the green form or photobleaching before detection in the red form. Further improvements in ELE may be achievable through additional optimization of sample preparation, imaging parameters and data processing. For instance, tuning laser wavelengths and power densities could help reduce photobleaching rates and thereby increase the ELE. In addition, localization algorithms specifically designed for high localization densities^35^ could potentially recover at least some of the substantial fraction of FPs discarded by conventional algorithms because they are already activated at the start of data acquisition.

We anticipate that the findings reported here for mEos4b will extend to most, if not all, PCFPs used as fluorescent markers in single-molecule based super resolution approaches. Although oxygen removal appears to be a viable strategy only for fixed cells, we expect it to improve imaging performance in nanoscopy approaches beyond PALM, such as iPALM^36^ or Minflux.^37^

## ASSOCIATED CONTENT

### Supporting Information

Materials and Methods (Generation and characterization of Nup96-mEos4b cells by CRISPR-Cas9; Sample preparation; PALM image acquisition; PALM data analysis; Oxygen level measurements; Statistical analysis; NMR experiments; SMIS simulations), Supporting Text,ReferencesFigures S1™S12; This material is available free of charge at https://pubs.acs.org/doi/xxxx.

## AUTHOR INFORMATION

### Authors

Oarii Vahirua − Univ. Grenoble Alpes, CEA, CNRS, Institut de Biologie Structurale (IBS), 38000 Grenoble, France;

Pascale Tacnet − Univ. Grenoble Alpes, CEA, CNRS, Institut de Biologie Structurale (IBS), 38000 Grenoble, France;

Jérémy Néri − Univ. Grenoble Alpes, CEA, CNRS, Institut de Biologie Structurale (IBS), 38000 Grenoble, France;

Anna Rosenkrancova − Univ. Grenoble Alpes, CEA, CNRS, Institut de Biologie Structurale (IBS), 38000 Grenoble, France;

Jip Wulffelé − Univ. Grenoble Alpes, CEA, CNRS, Institut de Biologie Structurale (IBS), 38000 Grenoble, France;

Oleksandr Glushonkov − Univ. Grenoble Alpes, CEA, CNRS, Institut de Biologie Structurale (IBS), 38000 Grenoble, France;

Virgile Adam − Univ. Grenoble Alpes, CEA, CNRS, Institut de Biologie Structurale (IBS), 38000 Grenoble, France;

Philippe Frachet − Univ. Grenoble Alpes, CEA, CNRS, Institut de Biologie Structurale (IBS), 38000 Grenoble, France;

Bernhard Brutscher − Univ. Grenoble Alpes, CEA, CNRS, Institut de Biologie Structurale (IBS), 38000 Grenoble, France ;

### Funding Sources

Financial support by the Agence Nationale de la Recherche (grants no. ANR-20-CE11-0013-01, ANR-22-CE13-0046 and ANR-22-CE11-0011-01), and the IR INFRANALYTICS (FR2054) is acknowledged. This work used the platforms of the Grenoble Instruct-ERIC center (ISBG; UAR 3518 CNRS-CEA-UGA-EMBL) within the Grenoble Partnership for Structural Biology (PSB), supported by FRISBI (ANR-10-INBS-0005-02) and GRAL, financed within the University Grenoble Alpes graduate school (Ecoles Universitaires de Recherche) CBH-EUR-GS (ANR-17-EURE-0003).

IBS acknowledges integration into the Interdisciplinary Research Institute of Grenoble (IRIG, CEA).

### Notes

The authors declare no competing financial interest.

## ACKNOWLEDGMENT

We thank Isabel Ayala and Salvatore de Bonis for help in protein purification, Jean-Baptiste Sibarita and Matthieu Sainlos for providing Dock-inert, Dock-catcher and mEos4b-tag plasmids, and Lydie Martin for initial attempts of sample deoxygenation using a glovebox.

## ABBREVIATIONS

FP: Fluorescent protein
PCFP: photoconvertible fluorescent protein
NPC: nuclear pore complex
LP: localization precision
ELE: effective labeling efficiency
PALM: photoactivated localization microscopy
MinFlux: Minimal Fluorescence Photon Fluxes Microscopy
iPALM: interference PALM
FRC: Fourier ring correlation
NMR: nuclear magnetic resonance
U2OS: human bone osteosarcoma
GLOX: glucose, glucose oxidase and catalase
HCS: High-content screening
ROS: reactive oxygen species

