## Supporting Information for "Optimizing environmental conditions for PALM imaging of the nuclear pore complex labeled with the photoconvertible fluorescent protein mEos4b"

### Materials and methods

#### Generation and characterization of Nup96-mEos4b cells by CRISPR-Cas9

##### *CRISPR-Cas9 plasmid construction*

Endogenous tagging of Nup96 with the photoconvertible fluorescent protein mEos4b was performed using a CRISPR-Cas9 paired-nickase strategy adapted from Koch *et al.*[1] targeting the Nup96-encoding region of the human *NUP98* gene. Two single-guide RNAs (sgRNAs), designated A12 (5′-GCAGTTCTCGCAGATAGGACT-3′) and B1 (5′-GTTGGGAGCCTGTGAGCCCC-3′), were designed to target opposite DNA strands near the intended insertion site. Each sgRNA was cloned separately into pX335-U6-Chimeric_BB-CBh-hSpCas9n(D10A) (Addgene #42335), which encodes the D10A nickase variant of *Streptococcus pyogenes* Cas9 and drives sgRNA transcription from the U6 promoter. Complementary oligonucleotides encoding the guide sequences (Eurofins Genomics) were annealed and ligated into BbsI-HF-digested pX335 (New England Biolabs). Following bacterial transformation and ampicillin selection, six colonies per construct were selected. Plasmid DNA was purified by miniprep, and insertion of the expected guide sequences was verified by Sanger sequencing using a primer directed against the U6 promoter.

##### *Construction of the mEos4b donor plasmid*

An ampicillin-resistant, pEX-A258-derived donor plasmid lacking a mammalian promoter was used to introduce mEos4b at the endogenous Nup96 C-terminus by homology-directed repair (HDR). The donor contained 801-bp left and right genomic homology arms flanking the insertion cassette. Nup96 and the fluorescent protein were separated by a 48-bp linker (5′-ACTAGTCGACGGTACCGCGGGCCCGGGATCCACCGGCCGGTCGCCACC-3′). The mEos4b donor was constructed using NEBuilder HiFi DNA Assembly (New England Biolabs, NEB). A 4,156-bp backbone fragment containing the genomic homology arms and a 721-bp fragment containing mEos4b, amplified from mEos4b/pRSET-B, were generated independently using Q5 High-Fidelity DNA Polymerase (New England Biolabs). Primers incorporated overlaps compatible with HiFi assembly. The amplicons were separated on agarose gels, excised and purified. Both fragments were assembled with NEBuilder HiFi DNA Assembly Master Mix and the resulting DNA was transformed into chemically-competent Beta-10 *E. coli* cells (NEB) on ampicillin-containing LB agar. Putative recombinant colonies were screened by colony PCR using a primer within the left homology arm and another within mEos4b. Plasmids from selected colonies were prepared and subjected to sequencing. Seven clones gave the expected sequence in the regions examined and their plasmid preparations were pooled for subsequent genome-editing experiments.

##### *Cell culture and CRISPR-Cas9 transfection*

U2OS cells were seeded 24 h before transfection in glass-bottom six-well plates at $2 \times 10^5$ cells per well in 3 mL of complete DMEM and incubated at 37°C. For each well, cells were co-transfected with 0.75 µg of pX335-A12, 0.75 µg of pX335-B1 and 1.5 µg of the Nup96-mEos4b donor plasmid. Plasmid DNA was diluted in Opti-MEM and mixed with FuGENE HD (Promega) at a FuGENE HD/DNA ratio of 3.5 µL/µg. A 150-µL volume of the transfection mixture was then added to each well, and cells were returned to 37°C.

##### *Flow cytometry and clonal isolation*

Following transfection, surviving U2OS cells were allowed to recover and expand for 8 days before cell sorting. Cells were trypsinized, resuspended in DMEM supplemented with NEAA, GlutaMAX and penicillin-streptomycin without FBS, and passed through 20-µm filters. Green-fluorescent cells were identified, sorted and cloned using a TYTO sorter (Miltenyi Biotec) combined with a Dispencell (Seed Bioscience). Cell populations were defined using forward- and side-scatter gating followed by successive doublet-discrimination gates, and green fluorescence was recorded in the GFP channel.

Sorted cells were distributed into four 96-well plates to derive individual clones. Each well contained 100 µL of a 1:1 mixture of U2OS-conditioned medium and fresh complete medium.

###### *Genomic PCR and assessment of allelic status*

Individual clones were characterized by genomic PCR to distinguish unmodified and mEos4b-tagged Nup96 alleles. PCR was performed using the forward primer 5'-GCCTTACCCAGTCCTATCTG-3′ and the reverse primer 5′-CAGAAGCCCTTATGCTACGTTCAGTATTAAG-3′. The expected amplification products were 236 bp for the unmodified locus and 974 bp for the mEos4b-tagged locus. PCR products were resolved by agarose-gel electrophoresis. Based on their amplification profiles, monoallelic clones displayed both the bands, whereas biallelic clones displayed only the mEos4b-tagged product and were selected for subsequent experiments.

##### Sample preparation

###### *Coverslip cleaning and preparation*

High-precision no. 1.5H glass coverslips (Marienfeld Superior, 0107052; 22 × 22 mm; thickness 170 ± 5 µm) were treated in a UV–ozone cleaner (UVOCS ®) for 20 min on each side. Subsequent handling was performed in a class II biological safety cabinet. Each coverslip was individually immersed in 70% (v/v) ethanol for 30 s with gentle agitation, transferred to a separate well of a six-well culture plate (Costar 3516, Corning; supplied by Dutscher), and allowed to dry for 1 h in the cabinet. Once dry, the coverslips were covered with 1 mL of complete culture medium per well and incubated for at least 24 h before cell seeding.

###### *Cell culture and seeding*

U2OS cells expressing Nup96–mEos4b or Nup96–mMaple (ref 300461, Cytion, Germany) were maintained in Dulbecco's modified Eagle medium (DMEM; Gibco, 11880028) supplemented with 1× MEM non-essential amino acids (Gibco, 11140-035), 1× GlutaMAX (Gibco, 35050-038), and 10% (v/v) fetal bovine serum (Gibco, 10270-106). Each prepared coverslip was seeded with $3 \times 10^5$ cells in complete culture medium and incubated at 37 °C in a humidified atmosphere containing 5% $CO_2$ for 24 h before fixation.

###### *Cell fixation and permeabilization*

The fixation sequence was adapted from the protocol for mMaple-tagged cells described by Thevathasan *et al.*[2] Coverslips were washed twice for 2 min in phosphate-buffered saline (PBS) and prefixed for 30 s in 2.4% (w/v) paraformaldehyde (PFA) in PBS, prepared by dilution of a methanol-free 16% aqueous stock solution (MP Biomedicals, 0219998380). After two PBS washes, cells were permeabilized in PBS containing 0.4% (v/v) Triton X-100 and 1% (w/v) bovine serum albumin (BSA) for 3 min. The coverslips were washed again in PBS and incubated in 2.4% (w/v) PFA in PBS for 30 min to complete fixation. After a PBS wash, residual aldehydes were quenched by incubation in 100 mM ammonium chloride ($NH_4Cl$) in PBS for 5 min. Coverslips were then washed twice in PBS and stored immersed in PBS at 4 °C until imaging.

###### *Live-cell preparation*

Immediately before live-cell imaging, cell-bearing coverslips were gently washed with phosphate-buffered saline (PBS) to remove residual culture medium. The coverslips were then mounted without fixation in the relevant imaging buffer using the slide–coverslip assembly described below.

###### *Sample mounting*

For imaging, 100 µL of the appropriate imaging buffer was dispensed into the cavity of a single-cavity microscope slide (Dutscher, 781645B; 76 × 26 mm; slide thickness 1.2–1.5 mm). The cell-bearing coverslip was placed over the cavity with the cells facing the buffer, and the perimeter of the slide–coverslip assembly was sealed with Twinsil Speed silicone (Picodent).

##### *In vitro sample preparation*

In vitro mEos4b immobilization employed the Dock-catcher/Dock-inert strategy based on the SpyTag/SpyCatcher technology,[3] described in ref 4. Initial attempts to collect data in the presence of GLOX to remove oxygen failed, producing overcrowded patterns. The reason for this behavior remains to be determined. Using $Na_2SO_3$ instead worked successfully.

##### *Imaging buffer preparation*

Imaging buffers were prepared immediately before use from 15× aqueous stock solutions. For a standard slide, 105 µL of working buffer was prepared by combining 98 µL of $H_2O$ or $D_2O$ with 7 µL of the appropriate 15× stock; 100 µL was then transferred to the slide cavity for mounting. $D_2O$-based buffers were prepared using $D_2O$ (Silantes GmbH, 300101400; isotopic enrichment >99.85 atom% D). The compositions below are final working concentrations. Values reported for $D_2O$-based formulations are nominal pH values.

**Table 1. Base imaging-buffer formulations.**

| Buffer | Final composition | Nominal pH | Solvent |
|---|---|---|---|
| TRIS | 50 mM TRIS; 140 mM NaCl | 8.0 | $H_2O$ or $D_2O$ |
| HEPES | 50 mM HEPES; 140 mM NaCl | 7.5 | $H_2O$ or $D_2O$ |
| Universal buffer | 50 mM sodium citrate; 50 mM glycine; 50 mM $KH_2PO_4$; 50 mM TRIS; 100 mM NaCl | 6.5, 7.5, or 8.5 | $D_2O$ |

For oxygen-depleted conditions, sodium sulfite ($Na_2SO_3$) was added from a 1 M stock to a final concentration of 47.6 mM, or a glucose oxidase/catalase mixture (GLOX/CAT) was used. The 10× GLOX/CAT stock contained 100 mM TRIS-HCl (pH 7.5), 25 mM KCl, 1 mM TCEP, 5 mg $mL^{-1}$ glucose oxidase (Sigma-Aldrich, G2133-50KU), 0.4 mg $mL^{-1}$ catalase (Sigma-Aldrich, C40), and 50% (v/v) glycerol. After centrifugation for 10 min at 10,000 × g, the stock was divided into 150 µL aliquots and stored at −20 °C. For a 150 µL GLOX/CAT working solution, 135 µL of 15% (w/v) glucose in $D_2O$, 10 µL of 15× HEPES stock, and 5 µL of 10× GLOX/CAT stock were combined.

Where indicated, a 100 mM Trolox stock was added to 1 mM final concentration. The stock was prepared by alkaline solubilization and characterized by absorbance at 255 nm; the batches used here contained approximately 12% Trolox-quinone (TQ), consistent with the redox-active Trolox/TQ system described by Cordes et al (ref 5 ). In the GLOX/CAT–Trolox formulation, 133.5 µL of 15% glucose in $D_2O$, 1.5 µL of 100 mM Trolox/TQ stock, 10 µL of 15× HEPES stock, and 5 µL of 10× GLOX/CAT stock were combined (150 µL final volume; 13.35% [w/v] glucose).

**Table 2. Imaging-buffer conditions used for PALM measurements.**

| Oxygen-scavenger | Base buffer | Solvent and nominal pH | Additive |
|---|---|---|---|
| None | TRIS | $H_2O$ or $D_2O$; pH 8.0 | None |
| None | HEPES | $H_2O$ or $D_2O$; pH 7.5 | None |
| None | Universal buffer | $D_2O$; pH 6.5, 7.5, or 8.5 | None |
| GLOX/CAT | HEPES | $D_2O$; pH 7.5 | None |
| $Na_2SO_3$ | HEPES | $D_2O$; pH 7.5 | None |
| $Na_2SO_3$ | Universal buffer | $D_2O$; pH 7.5 | None |
| GLOX/CAT | HEPES | $D_2O$; pH 7.5 | 1 mM Trolox/TQ |
| $Na_2SO_3$ | HEPES | $D_2O$; pH 7.5 | 1 mM Trolox/TQ |
| $Na_2SO_3$ | Universal buffer | $D_2O$; pH 7.5 | 1 mM Trolox/TQ |

*Note: Trolox concentration refers to total Trolox derived species in the characterized TX/TQ stock.*

#### PALM image acquisition

PALM data were collected on a custom-built PALM/STORM microscope based on an Olympus IX81 inverted microscope equipped with a 100× oil-immersion apochromatic objective with a numerical aperture of 1.49 (Olympus). The setup incorporated 405 nm (CrystaLaser) and 561 nm (Cobolt) lasers and fluorescence images were recorded with an ORCA-Fusion BT scientific (sCMOS) camera (Hamamatsu). For a detailed description, see ref [6].

mEos4b red fluorescence was collected through an RFP filter cube containing a 580 nm-edge dichroic beamsplitter (Semrock, FF580-FDi01-25x36) and a 630/92 nm emission bandpass filter (Semrock, FF01-612/69-25) in front the camera.

Data acquisition was performed with Micro-Manager 2.0.1and laser illumination sequences were generated with Labview 2025. 405 nm illumination was used to photoconvert the fluorescent proteins, and 561 nm illumination was used to excite their red state. The 561 nm irradiance at the sample was ~1 kW $cm^{-2}$. The 405 nm irradiance was increased progressively according to a quadratic ramp, starting from 0 and reaching a maximum of ~10 W $cm^{-2}$. Each illumination cycle comprised 80 ms of 561 nm illumination, 5 ms without illumination, 5 ms of 405 nm illumination, and a further 10 ms without illumination, giving a total cycle duration of 100 ms. The camera exposure time was 80 ms, and 20,000 frames were recorded per field of view. In all conditions, only very few localizations per frame were still present at the end of the acquisitions.

For pH-dependent measurements with universal buffer (Fig. S4), no laser power calibration was done. As a consequence, the extracted absolute Nphot values cannot be reliably compared to those of other experiments and only relative values are show in Fig. S4.

For live-cell experiments, the illumination sequence was adapted to a total cycle duration of 80 ms. Each cycle comprised 50 ms of 561 nm illumination, 10 ms without illumination, 5 ms of 405 nm illumination, and a further 15 ms without illumination. The camera exposure time was 50 ms, and 10,000 frames were recorded per field of view.

For in vitro experiments, the illumination sequence was similar to that employed for fixed-cell experiments, except that the power density of the 405 nm laser was kept constant throughout the acquisitions at ~2 W/$cm^2$. Stacks consisted of only 10,000 frames.

#### Oxygen level measurements

Oxygen in the imaging chamber formed by a cavity slide and a coverslip was measured using a PICO-$O_2$ optical oxymeter supplied by Bionef (P-PICO-O2) and trace-range self-adhesive oxygen sensor spots (P-TROXSP5-ADH). The instrument was calibrated according to the manufacturer's instructions, using a P-OXCAL capsule for the zero-oxygen calibration point. A sensor spot was attached to the bottom of the cavity containing the imaging buffer. A volume of 100 µL of buffer was dispensed into the cavity, which was covered with a no. 1.5H coverslip and sealed with Twinsil Speed silicone (Picodent), following the same procedure as for microscopy samples. Oxygen was measured at successive time points through the glass.

The measurements, recorded as an equivalent oxygen partial pressure in hPa, were converted to dissolved oxygen concentration in µmol·L$^{-1}$ using Henry's law:

$$C(\mu\text{mol} \cdot \text{L}^{-1}) = 10^6 \, \kappa_\text{H} \, \frac{p_{\text{O}_2}(\text{hPa})}{1013.25} \quad (1)$$

The measurements were performed at room temperature, approximately 20 °C, and the calculations used the nominal coefficient $\kappa_\text{H} = 1.28 \times 10^{-3}$ mol·L$^{-1}$·atm$^{-1}$, giving C [µMol] ≈ 1.2633 × $p_{\text{O}_2}$ [hPa]. This conversion provides an estimate because the coefficient depends on temperature and medium composition and was not determined specifically for each imaging buffer. Typical measurements are reported in Fig. S12.

#### PALM data analysis

Localization tables (.csv) were used to characterize effective labelling efficiency (ELE), photons per localization (Nphot), localization precision (LP) and image resolution based on Fourier Ring Correlation (FRC). NPC analysis followed the general scheme presented in Thevathasan *et al.*[2] but was based on custom scripts written in Matlab 2025b.

##### *Localization data and preprocessing*

PALM image sequences were analyzed in ThunderSTORM.[7] An initial drift correction was performed in ThunderSTORM by using the cross-correlation method. Localizations with *uncertainty_xy* > 12 nm and *sigma* < 80 or > 145 nm were then filtered out. To remove isolated localizations, a 2D density filter was applied with a neighborhood radius of 50 nm and a minimum-neighbor setting of 5. The exported coordinates were subsequently subjected to an additional drift correction with COMET.[8]

##### *Photons per localization and localization precision*

Nphot and LP (*intensity* and *uncertainty_xy* in ThunderSTORM) were calculated for each field of view (nucleus) as median values from the selected filtered localizations as described above.

##### *Effective Labelling Efficiency*

For each Field of View (FOV), NPCs were searched within manually selected nuclear regions of interest. To segment NPCs, localizations were rendered on a 10×10 nm² grid and convolved with a disk of radius 55 nm. Candidate NPCs were first detected by a circular Hough transform. Candidates with at least 20 localizations were selected. A circle was then fitted from those localizations to refine NPCs centers and radii. The median fitted radius defined the effective NPC radius $r_0$ for each FOV. NPCs with radii out of bounds set to $r_0 \pm 2\times lp_0$ where $lp_0$ is the median localization uncertainty over each FOV were rejected. A counting annulus with bounds $r_0 \pm 3\times lp_0$ was subsequently defined. NPCs that contained more than 30% or 70% of localizations out of this annulus, (inside or outside respectively), were further rejected. Corner counting used localizations within the annulus. Localization coordinates were converted to polar coordinates around $r_0$, rotated to align their eightfold angular pattern and divided into eight 45° sectors. A sector containing at least one localization was considered a visible corner. Each pore therefore contributed a corner count k from 0 to 8. The ELE parameter was then calculated using a binomial distribution model as described in Thevathasan *et al*,[2] with ELE uncertainties estimated by bootstrapping.

##### *Image resolution by Fourier Ring Correlation*

Image resolution was estimated by Fourier ring correlation (FRC) using a custom Python application following established procedures for localization microscopy.[9] Localizations were split according to odd and even acquisition frames and reconstructed independently on a 5 nm pixel grid. Reconstructions were Gaussian-smoothed ($\sigma$ = 5 nm) and an inward boundary taper of two pixels was applied before Fourier transformation. FRC curves were smoothed using a centered five-bin moving average, and resolution was determined from the first downward crossing of the 1/7 threshold. Cutoffs $\geq$ 0.49 cycles per pixel were considered Nyquist limited, corresponding to a resolution of approximately 10.2 nm for a 5 nm reconstruction pixel size.

##### *Analysis of in vitro data*

*In vitro* data were analyzed in ThunderSTORM.[7] Fluorescence traces from individual mEos4b molecules were then reconstituted based on in-house developed Matlab scripts, as described in ref [10], from which the different photophysical parameters of Fig. S9 were extracted. To calculate the number of photons emitted per localization before photobleaching, only traces comprising at least two localizations were considered, and the last localization from those traces (during which irreversible bleaching occurred) was omitted in the calculation.

#### Statistical analysis

Each independently prepared coverslip was considered one independent sample. For each FOV (typically a nucleus) acquired on that coverslip, the Nphot and LP median values were calculated. Arithmetic means of these values were then obtained from all analyzed FOVs from a single coverslip (typically 3). Reported sample sizes in Figures denote the number of independently prepared coverslips, except for **Fig.** S3 and S9 in which sample sizes correspond to the total number of FOVs.

Statistical analyses were performed in GraphPad Prism 10. Two independent groups were compared using the Mann–Whitney U test. Comparisons involving more than two independent groups were evaluated with the Kruskal–Wallis test. Tests were two-tailed, with a significance threshold of $\alpha$ = 0.05. Exact p values and the number of independent measurements are reported in the corresponding figure legends.

#### NMR experiements

##### *Sample Preparation*

mEos4b was recombinantly expressed in Escherichia coli  and uniformly 13C/15N-labeled using standard procedures. The labeled protein was purified using Ni-NTA affinity chromatography, followed by size-exclusion chromatography on a Superdex 75 column. NMR samples were prepared at a protein concentration of 460 µM in 20 mM HEPES buffer (pH 7.5) and transferred into standard 3 mm NMR sample tubes. To prepare oxygen-depleted and oxygen-enriched samples, the solutions were bubbled with argon or oxygen, respectively, for approximately 10 min directly in the NMR tubes, which were subsequently sealed.

##### *NMR Spectroscopy*

NMR photoconversion experiments were performed at 35°C on a Bruker Avance NEO spectrometer operating at a $^{1}$H Larmor frequency of 950 MHz. The spectrometer is equipped with a 5-mm helium-cooled triple-resonance probe (HCN) and pulsed z-field gradients. To monitor the photoconversion, a series of 2D $^{1}$H-$^{15}$N BEST-TROSY correlation spectra[11] were recorded with an acquisition time of 11.4 min per experiment, and an overall experimental time of 23.5 hours. Sample illumination was achieved using an inversed double tube setup, adapted from ref [12], with the NMR sample in the inner

(3 mm) tube, and the outer D2O-filled 5 mm tube acting as light guide. The sample was continuously illuminated in situ in the NMR spectrometer using a 405 nm laser source at a power of about 3.4 mW/cm$^2$ (fibre output). NMR data were processed using TopSpin v4.1 and CCPNMR v3 software.[13]

##### *Data Analysis*

To quantify the difference in photoconversion yield between oxygen-depleted and oxygen-enriched samples, the NMR kinetics of 16 well-resolved, non-overlapping red-state peaks were analysed. To account for small differences in protein concentration between the samples, the green-state NMR signal intensities were measured before illumination and compared. The resulting red-state kinetic traces were then normalized using a correction factor of 0.95 to account for the difference in initial green-state signal intensity. This normalization enables the red-state NMR intensities to be compared directly between samples, thereby reflecting differences in red-state population. A selection of 4 (out of 16) normalized kinetic traces are shown in **Fig.** 3. The relative increase in maximum red-state population achieved was then determined from the maximum intensity in the oxygen-depleted and oxygen-enriched samples.

Additional NMR data fitting to a photophysical model was performed, as shown in Fig. S11. Gly31 was selected for detailed kinetic analysis because all four states (GA, GB, R1, and R2) give rise to well-resolved NMR signals. As GA and GB undergo chemical exchange on the minute timescale and exhibit indistinguishable photoconversion kinetics, their individual signal intensities were summed to represent the overall green-state population (G). The state transitions were described by a system of first-order linear differential equations based on the kinetic model shown in Fig. S11A and implemented in a custom Python script. To quantify kinetic differences between oxygen-enriched and oxygen-depleted conditions, the two recorded datasets were fitted simultaneously. The photoconversion rate constant ($k_{G\rightarrow R1}$), the red-state interconversion rate constants ($k_{R_1\rightarrow R_2}$, $k_{R_2\rightarrow R_1}$), and the red-state photobleaching rate constant ($k_{R_1\rightarrow B_R}$) were treated as global parameters shared between both datasets. Of note, when allowed to vary independently, the photoconversion rate constants $k_{G\rightarrow R1}$ yielded indistinguishable values between the two datasets within experimental error, indicating no significant oxygen dependence. The red-state photobleaching rate was also kept global because the slow decay of the red plateau is too noisy to reliably resolve oxygen-dependent differences. In contrast, the green-state photobleaching rate constant ($k_{G\rightarrow B_G}$) was allowed to vary independently between the two conditions, as its oxygen dependence is determined by the distinct red-state plateau height difference between the two samples.

#### SMIS simulations

Simulations were performed using the SMIS software,[14] using the photophysical model presented in Fig. S10 and the same main imaging parameters (number of frames, frame time, pixel size, laser wavelengths and power densities, emission filter) as employed in real experiments except that the simulated detector was an EMCCD. A first pair of simulations (in the presence and absence of oxygen) aimed to reproduce the *in vitro* mEos4b data, with sparsely distributed molecules and a fixed 405 nm power density over the whole acquisition. These simulations allowed to reconstruct full fluorescence traces, so that the results can be compared to those of Fig. S9. To simulate the GLOX condition, all the oxygen-dependent pathways in the model of Fig. S10 (oxidation of $T_1$ to a cationic radical and direct photobleaching from $T_1$) were shut off, and the lifetime of $T_1$ was increased by 20%. The complete list of employed photophysical parameters is presented in Table S1. The second pair of simulations aimed to reproduce the NPC data, notably the increase in ELE observed in the absence of oxygen. 200 NPCs per field of view were simulated, assuming 32 mEos4b molecules per NPC, a maturation efficiency of 90%, and 2 molecules per corner. It is important to consider the results of the simulations only as qualitative, as the photo transformation quantum yields and rates in Table S1 are guessed parameters guided by literature values.

**Supporting Table 1**: *Phototransformation quantum yields and thermal rates of the mEos4b photophysical model used in SMIS simulations.*

| | ***Green anionic*** | ***Green neutral*** | ***Green T₁*** | ***Green dark anionic*** | ***Green dark cationic*** | ***Green bleached*** | ***Red anionic*** | ***Red neutral*** | ***Red T₁*** | ***Red dark anionic*** | ***Red dark cationic*** | ***Red long-lived dark*** | ***Red bleached*** |
|---|---|---|---|---|---|---|---|---|---|---|---|---|---|
| ***Green anionic*** | | Rapid Exchange | $1\times10^{-3}$ | 0 | 0 | $2.5\times10^{-8}$ | 0 | 0 | 0 | 0 | 0 | 0 | 0 |
| ***Green neutral*** | Rapid Exchange | | $1\times10^{-3}$ | 0 | 0 | $2.5\times10^{-8}$ | 0 | $2.0\times10^{-4}$ | 0 | 0 | 0 | 0 | 0 |
| ***Green T₁*** | 0 | $(1000\ s^{-1})$* $(800\ s^{-1})$* | | $(50\ s^{-1})$* | $(50\ s^{-1})$* $(0\ s^{-1})$* | $5\times10^{-6}$ $(5\ s^{-1})$* $(1\ s^{-1})$* | 0 | 0 | 0 | 0 | 0 | 0 | 0 |
| ***Green dark anionic*** | 0 | $3\times10^{-3}$ $(20\ s^{-1})$* | | | 0 | $1\times10^{-6}$ $(0.25\ s^{-1})$* | 0 | 0 | 0 | 0 | 0 | 0 | 0 |
| ***Green dark cationic*** | 0 | $3\times10^{-3}$ $(20\ s^{-1})$ | | 0 | | $1\times10^{-6}$ $(0.25\ s^{-1})$* | 0 | 0 | 0 | 0 | 0 | 0 | 0 |
| ***Red anionic*** | | | | | | | | Rapid Exchange | $1.0\times10^{-3}$ | 0 | 0 | $4.5\times10^{-5}$ | $1.0\times10^{-7}$ |
| ***Red neutral*** | | | | | | | Rapid Exchange | | $1.0\times10^{-3}$ | 0 | 0 | $4.5\times10^{-5}$ | $1.0\times10^{-7}$ |
| ***Red T₁*** | | | | | | | $5.0\times10^{-4}$ $(1000\ s^{-1})$* $(800\ s^{-1})$* | 0 | | $(50\ s^{-1})$* | $(50\ s^{-1})$* $(0\ s^{-1})$* | 0 | $1.3\times10^{-5}$ $(5\ s^{-1})$* $(2\ s^{-1})$* |
| ***Red dark anionic*** | | | | | | | $3.0\times10^{-3}$ $(20\ s^{-1})$ | 0 | 0 | | 0 | 0 | $1.3\times10^{-6}$ $(0.25\ s^{-1})$* |
| ***Red dark cationic*** | | | | | | | $3.0\times10^{-3}$ $(20\ s^{-1})$ | 0 | 0 | 0 | | 0 | $1.3\times10^{-6}$ $(0.25\ s^{-1})$* |
| ***Red long-lived dark*** | | | | | | | $3.0\times10^{-2}$ $(1.5\times10^{-3}\ s^{-1})$ | 0 | 0 | 0 | 0 | | $2.0\times10^{-7}$ |

* Value in parentheses refer to thermally induced relaxation rates; $ Values in green and blue refer to rates that are different when $O_2$ is removed;

#### Supporting Text : relationship between ELE, LP, Nphot and FRC resolution

The four metrics reported in this work are interconnected.

LP is strongly correlated with Nphot through the Thompson formula.[15] However, as LP also depends on the standard deviation of background fluorescence, the correlation is not strict. This is exemplified in the situation reported in Fig. 2, where the addition of GLOX increases both Nphot and background fluorescence, and as a consequence does not substantially improve LP.

The ELE in principle should not correlate with LP or Nphot, but it should correlate with the FRC resolution. In practice, things are different when dealing with NPCs.

First, the analysis method to extract the ELE is based on counting localizations in each NPC corner (see Materials and Methods). Localizations with poorer precision may be found in corners they do not belong to, introducing errors in the extracted ELE. Typically, an overall decrease in LP results in more corner misassignment and ELE overestimation. Thus, extracted absolute ELE values should be considered with care, as they may depend on fluorophore choice, imaging parameters and processing pipeline, notably localization filtering. Only relative ELE values calculated on identically filtered data should be considered reliable. Thus in this work, we do not attempt to compare the obtained ELE's with previously published values, although our ELE's typically fall in the same range as those published in e.g. ref 2.

Concerning filtering, it is interesting to note that, for HEPES, pH7.5, $D_2O$ conditions, on average, ~45% of the initially selected NPCs in our processing pipeline are eventually rejected based on the criteria described above. This percentage drops to ~35% when GLOX is added. The reduced number of rejected NPCs in the absence of $O_2$ correlates with the observed increased image quality.

Second, while in theory a high ELE should correlate with a high FRC resolution due to better sampling of the biological structures (Nyquist sampling theorem), in the case of NPCs this is not necessarily observed. This is shown by SMIS simulations[14] in which we compare two data sets obtained in identical imaging conditions and with the same simplified mEos4b photophysical model (giving rise to 100% photoconversion), except for a difference in maturation level (20% to 50%).It can clearly be seen in Fig. S5 that, while the two datasets give rise to different ELE's, they give rise to the same FRC resolution. The reason behind this observation is likely the fact that the spatial frequency content of incomplete NPC rings is not much different than for complete rings. Nevertheless, the FRC resolution does correlate with LP, as accurately positioned fluorophores give rise to better defined NPC rings.

#### Supporting Figures

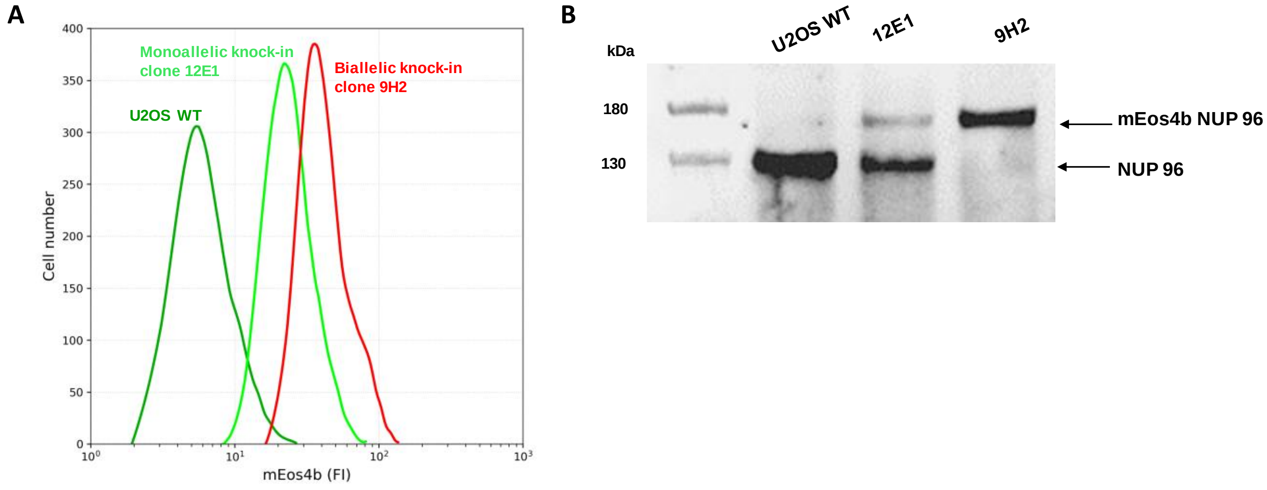

**Figure S1: Generation and characterization of CRISPR/Cas9-edited U2OS clones expressing mEos4b-tagged Nup96.** (A) Flow cytometry analysis of wild-type U2OS cells and selected CRISPR/Cas9-edited clones. (B) Western blot analysis of Nup96 and the Nup96-mEos4b fusion protein using a rabbit polyclonal antibody against Nup96 under reducing conditions following SDS–PAGE. The genomic status of the selected clones was verified by PCR, as described in the Materials and Methods. Clone 12E1 was identified as a monoallelic knock-in, whereas clone 9H2 was identified as a biallelic knock-in and selected for further imaging experiments. The expected molecular weights of Nup96 and the Nup96-mEos4b fusion protein are approximately 130 and 180 kDa, respectively.

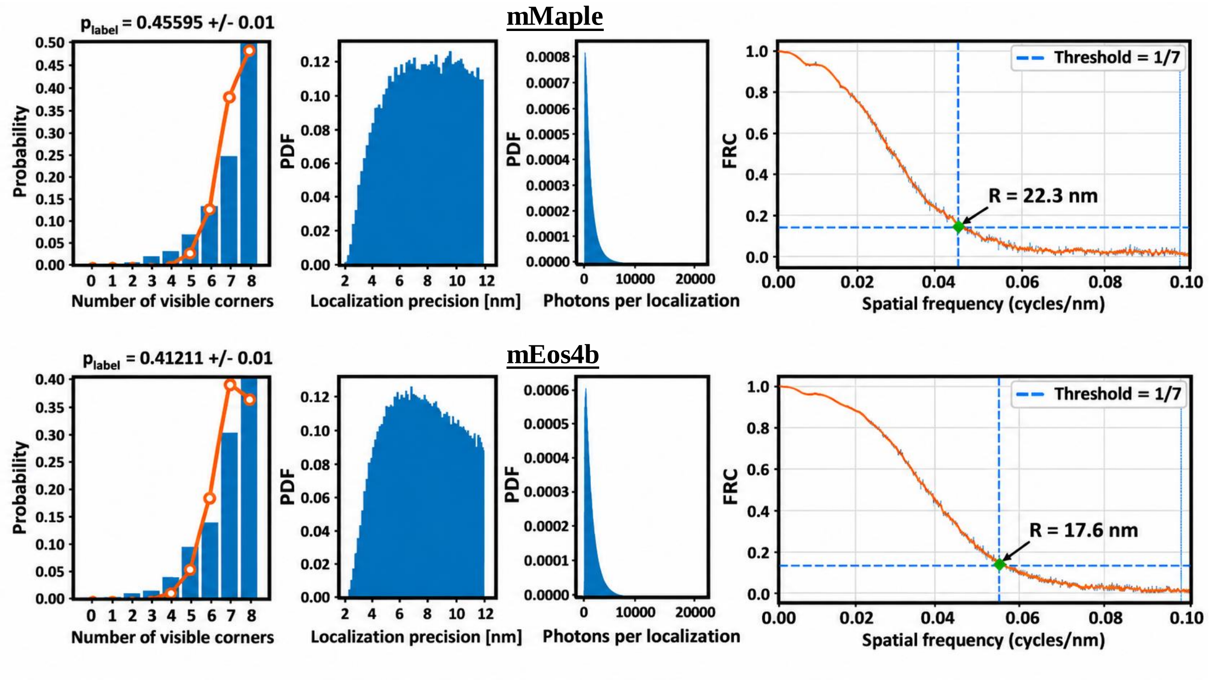

**Figure S2: Representative analysis of effective labeling efficiency, localization precision, number of photons per localization and FRC resolution for mMaple and mEos4b PALM imaging.**

Quantitative characterization of NPCs in fixed U2OS cells expressing Nup96-mMaple (top row) or Nup96-mEos4b (bottom row) under reference environmental conditions. Left: Probability distributions of the number of visible NPC corners (blue bars), ranging from 0 to 8, together with the corresponding model-fitted distributions (orange curves). The estimated effective labeling efficiency (ELE, expressed as $p_{label}$) is indicated above each graph. Middle left: Probability density functions (PDFs) of localization precision, showing the distributions of individual localization uncertainties for each fluorescent protein, after applied filtering (see Materials & Methods). Middle right: PDFs of photon numbers per localization after applied filtering. Right: Fourier ring correlation (FRC) curves obtained from independently reconstructed localization subsets, displaying the raw (blue) and smoothed (orange) correlation profiles as a function of spatial frequency. The horizontal dashed lines indicate the 1/7 FRC threshold, while the vertical dashed lines mark the corresponding cutoff frequencies used to determine the spatial resolution.

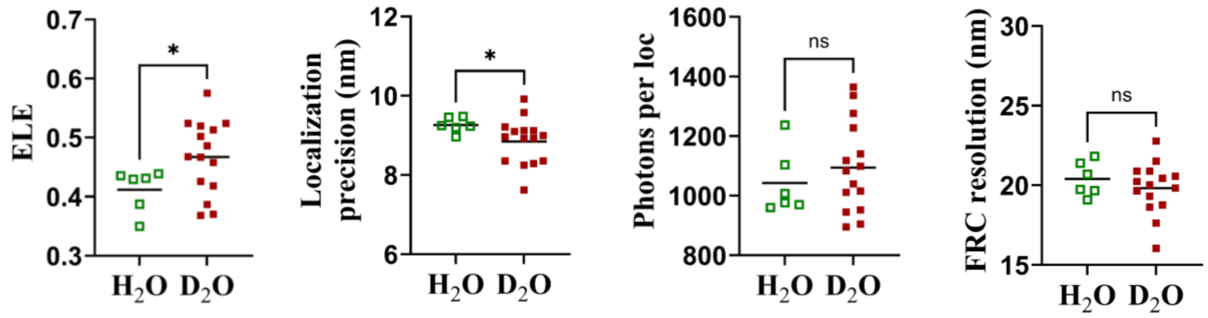

**Figure S3: Effect of solvent deuteration on mEos4b PALM imaging performance under oxygenated conditions.**

ELE, median localization precision, median photons detected per localization and Fourier ring correlation (FRC) resolution were compared between fixed U2OS Nup96–mEos4b cells imaged in $D_2O$- or $H_2O$-based buffers under otherwise equivalent oxygenated conditions. Individual symbols represent independent samples prepared in TRIS or HEPES buffers (pooling samples prepared in both buffers provide buffer-independent statistics). ($D_2O$, n = 15; $H_2O$, n = 6). Horizontal lines indicate group means. Pairwise comparisons were performed using a two-tailed unpaired Mann_Whitney U test. Statistical significance is indicated as : ns, not significant, *, $p < 0.05$

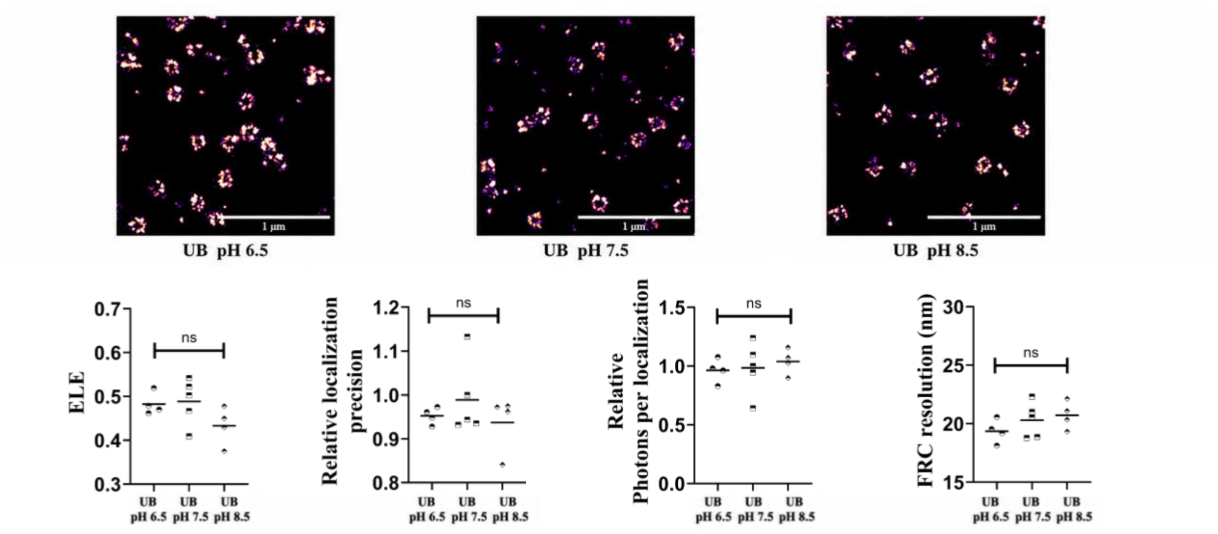

**Figure S4: Effect of pH on mEos4b PALM performance using universal buffer (UB).** Representative PALM reconstructions of fixed U2OS Nup96-mEos4b cells acquired in universal buffer (UB), in $D_2O$, adjusted to pH 6.5, pH 7.5, or pH 8.5. Scale bars, 1 µm. Quantitative comparisons of ELE, localization precision, photons per localization, and FRC resolution are shown below representative images. Each symbol represents one coverslip, with at least three spatially distinct fields of view (FOVs), analyzed and averaged to obtain a single coverslip-level value. Relative values are shown for localization precision and photons per localization. Horizontal bars indicate the mean. Statistical differences among pH conditions were assessed using Kruskal-Wallis tests followed by Dunn's multiple-comparisons pot hoc test. ns, not significant ($p > 0.05$).

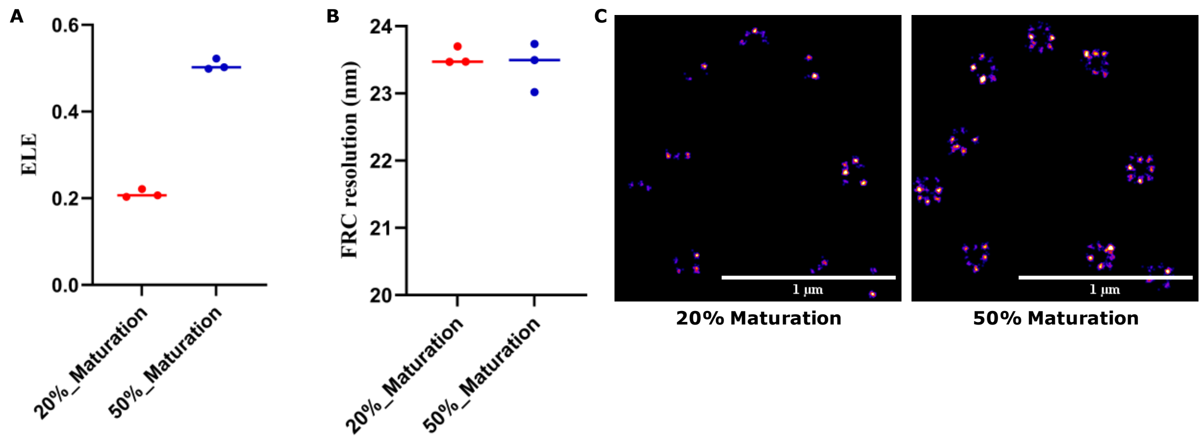

**Figure S5: In silico assessment of the relationship between ELE and FRC resolution for NPC imaging.**

Simulated NPC datasets were generated using fluorophore maturation fractions of 20% and 50% ($n$ = 3 independent simulations per conditions) to modulate the fraction of detectable Nup96-associated fluorophores and thereby the ELE. (A) ELE and (B) FRC resolution obtained for each simulated dataset. (C) Parts of reconstructed images from SMIS-generated data sets for illustrative purpose.

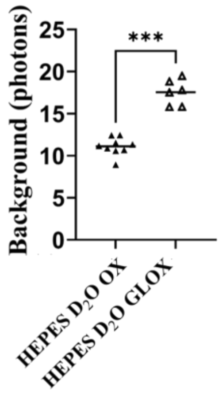

**Figure S6: Effect of GLOX on fluorescence background**

Quantitative comparisons of the standard deviation of fluorescence background for each localization in the presence or absence of GLOX. Representative images are shown on the right. For each independent experiment, at least three spatially distinct fields of view (FOVs) were acquired and analyzed. Each parameter was determined independently for each FOV, and the resulting values were averaged to obtain a single coverslip-level measurement. Each symbol represents one independent experiment (one coverslip), and horizontal lines indicate group means. Statistical comparisons were performed using two-tailed, unpaired Mann–Whitney U tests. Statistical significance is indicated as *** ($p < 0.001$).

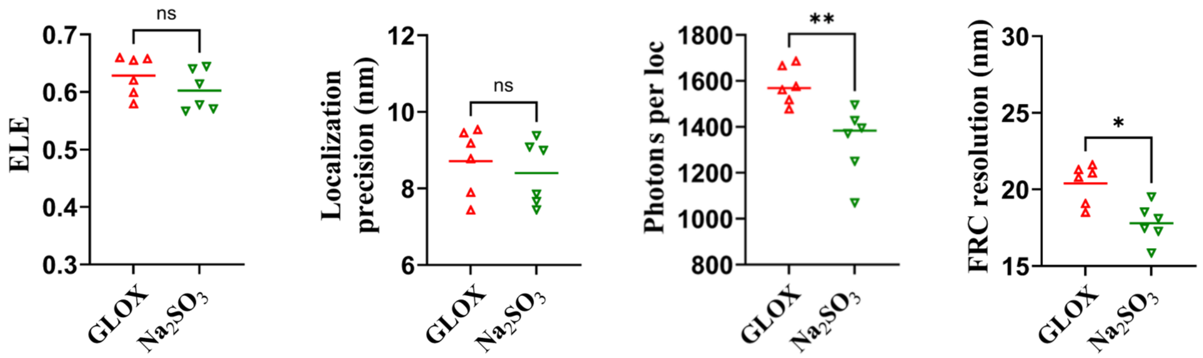

**Figure S7: Comparison of glucose oxidase/catalase and sodium-sulfite oxygen-scavenging systems on PALM performance.**

ELE, median localization precision, median number of photons per localization and FRC resolution were compared between samples imaged using a glucose oxidase/catalase (GLOX) or $Na_2SO_3$ under otherwise identical buffer conditions. Individual symbols represent independent measurements and horizontal bars indicate the group mean value. Pairwise comparisons were performed using two-tailed unpaired Mann-Whitney U tests. ns, not significant; *, $p < 0.05$; **, $p<0.001$.

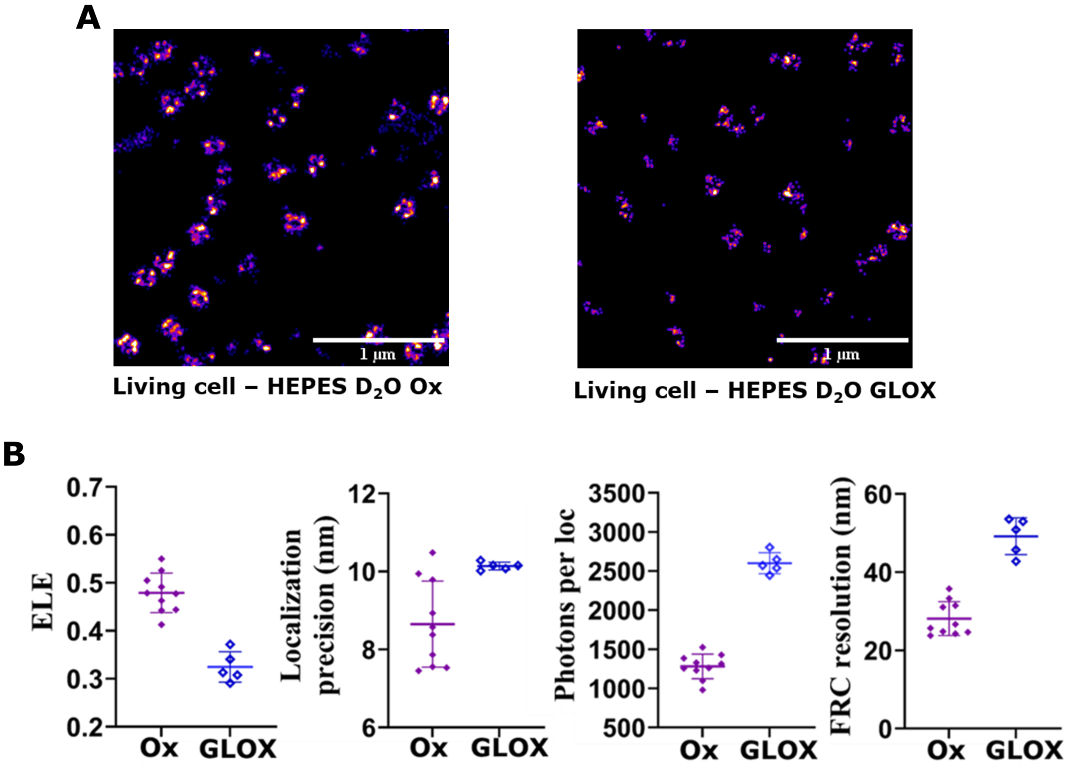

**Figure S8: mEos4b PALM imaging in living cells under oxygenated and oxygen-scavenging conditions.** (A) Representative PALM reconstructions of NPCs in living U2OS Nup96-mEos4b cells acquired in HEPES buffer prepared in $D_2O$ under oxygenated conditions (left) or supplemented with a glucose oxidase/catalase (GLOX) oxygen-scavenging system (right). Scale bars, 1 µm. (B) Quantitative comparison of ELE, median number of photons per localization, median localization precision, and FRC resolution between the two imaging conditions (from left to right). Each symbol represents one analyzed cell, and horizontal lines and error bars indicate group means and standard deviations (SD), respectively. Significance tests are not shown as many data sets recorded in the presence of GLOX could not be processed and were left over.

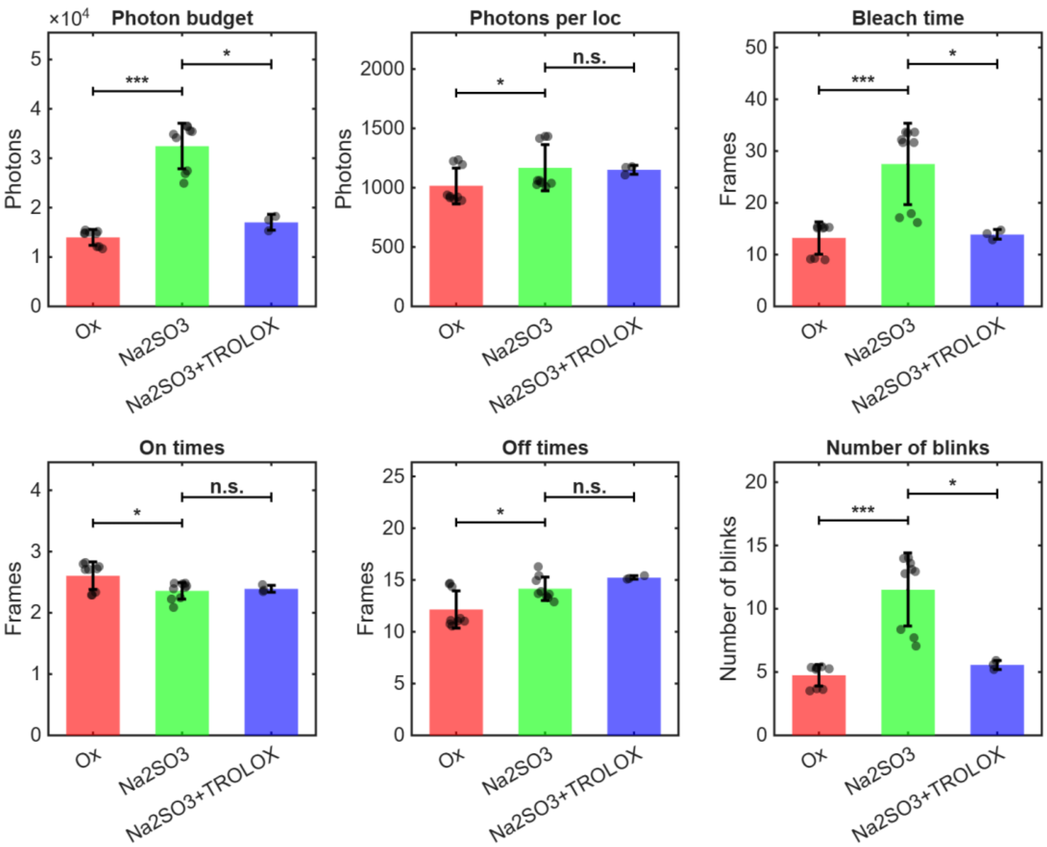

**Figure S9: Photophysical parameters of mEos4b immobilized in vitro in the presence or absence of oxygen.** Quantitative comparison of total photon budget, number of photons per localization (restricted to localizations free of photobleaching, see main text), bleach time, on and off times and number of blinks (median values) between oxygenated and deoxygenated conditions. Each symbol represents one field of view (FOV). 3 FOVs from 2 independent experiments were collected. Error bars indicate group standard deviations. Statistical differences were assessed using Kruskal-Wallis tests. *, $p < 0.05$, ***; $p < 0.001$; ns, not significant ($p > 0.05$).

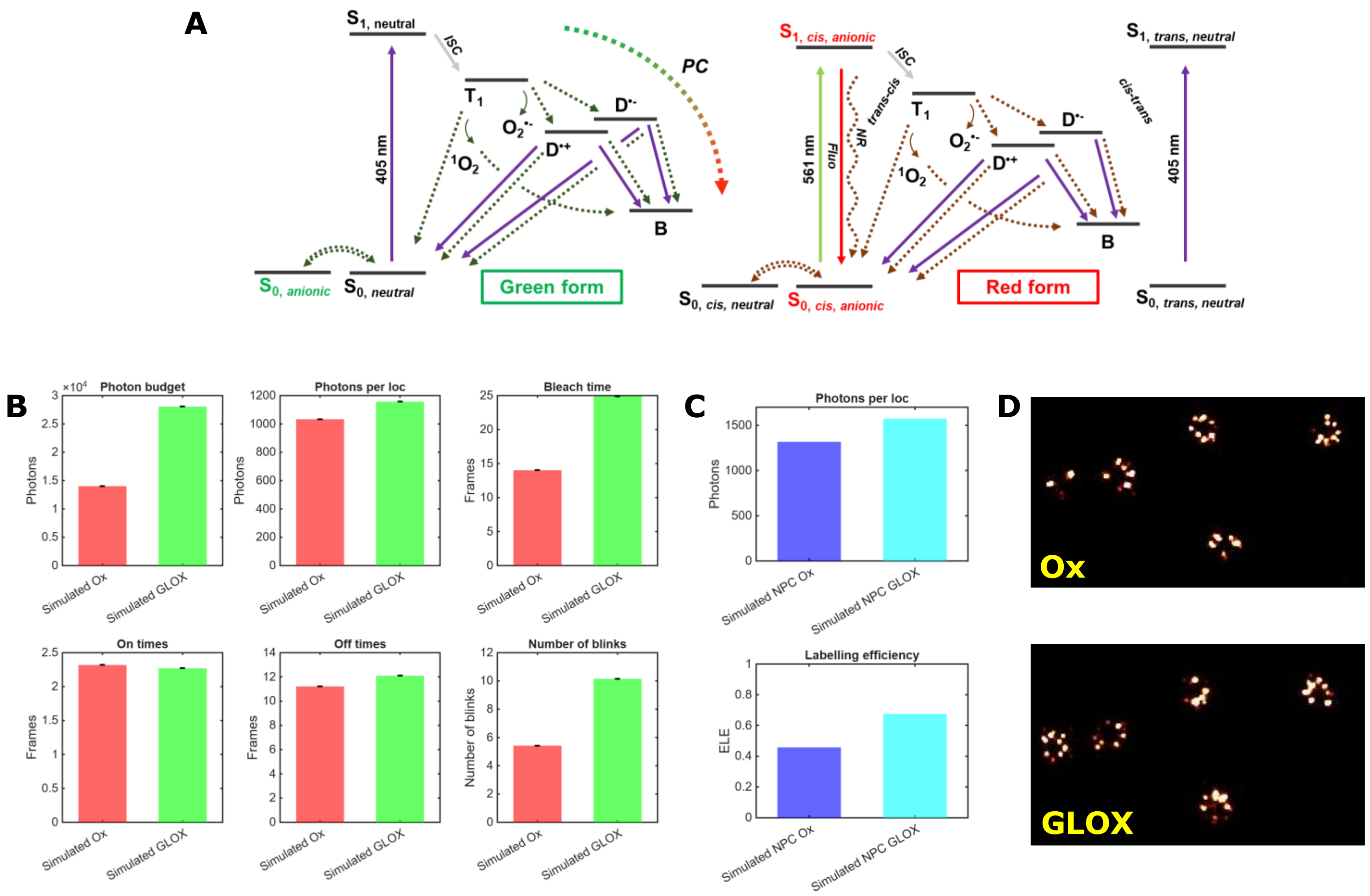

**Figure S10: Simulated data recapitulating experimental observations**

(A) Photophysical model used for SMIS simulations. As compared to the model of Fig. 4, additional, $O_2$-independant, sources of blinking of mEos4b (anionic radical in the green and red form and cis-trans isomerization in the red form) have been included. S: Singlet state, T1: Triplet state, B: Bleached, $D^{\bullet+}$: Cationic radical, $D^{\bullet-}$: Anionic radical, PC: photoconversion, Fluo: Fluorescence, NR: Non radiative relaxation. (B): Photophysical parameters extracted from SMIS *in vitro* (sparsely distributed mEos4b) simulated data in the presence (Ox) or absence (GLOX) of oxygen. (C): ELE and Nphot extracted from SMIS *in cellulo* (mEos4b-labeled NPCs) simulated data in the presence (OX) or absence (GLOX) of oxygen. No error bar is shown in (B) and (C) since simulated data are fully reproducible (D): Representative extracts of simulated PALM images of NPCs

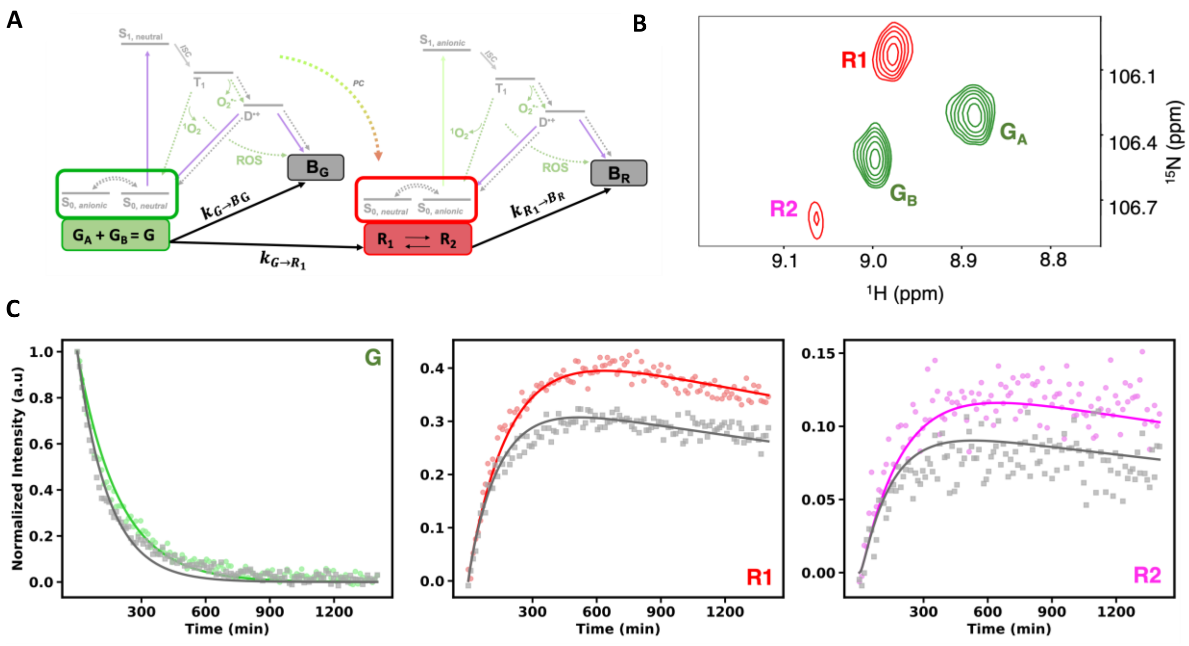

**Figure S11: NMR data fitting to photophysical model**. A: Photophysical model derived from Fig. 4, accounting for the long-lived species (GA, GB, R1, R2) observed by NMR, as well as the photobleached green (BG) and red (BR) species. For kinetic fitting, GA and GB were treated as a single green-state population, since exchange between them occurs on a much faster time scale (~ 1 min) than photoconversion **(**ref [16]**)**. In contrast, R1 and R2 were considered as distinct kinetic species (ref [6]). B: Spectral region of a 2D 1H-15N spectrum showing 4 correlation peaks, assigned to the GA, GB, R1 and R2 states of Gly-31. This residue was chosen for kinetic data fitting, because all 4 signals are spectrally resolved, allowing their NMR intensities to report on the relative populations of the corresponding states. C: Kinetic traces for the combined green state G = GA+GB, R1, and R2, extracted from two data sets acquired under low-power UV illumination for oxygen-rich (grey), and oxygen-depleted (colored) samples. Solid lines represent the results of a global fit of both data sets to the model in A. Only the bleaching rate constant of the green state ($k_{G\to B_G}$) was allowed to depend on oxygen availability and all other rate constants were shared between the two datasets. The fitting results suggest that under oxygen-depleted conditions, the green-state photobleaching rate decreases by approximately 44 % compared to the oxygen-enriched sample (from $(3.81 \pm 0.07) \times 10^{-3}\ \mathrm{min}^{-1}$ to $(2.15 \pm 0.05) \times 10^{-3}\ \mathrm{min}^{-1}$), suggesting that this mechanism is at the basis of the increased apparent photoconversion yield.

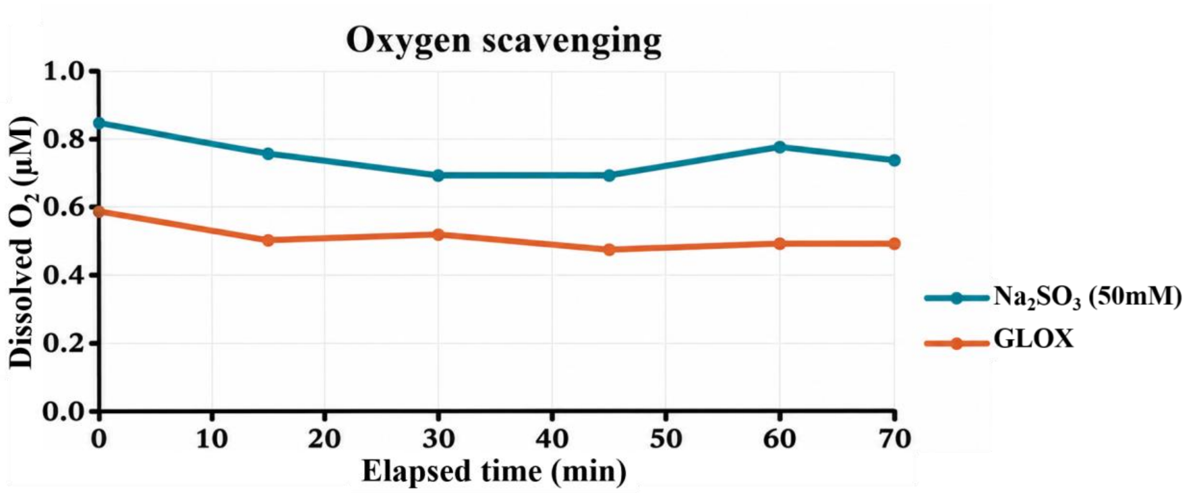

**Figure S12: Experimental characterization oxygen-scavenging systems used for PALM imaging.**

Evolution of dissolved $O_2$ concentration in samples containing GLOX or sodium sulfite ($Na_2SO_3$). Oxygen measurements were performed in sealed chambers to reproduce the conditions used during PALM acquisitions. Oxygen concentration in typical aerated solutions is ~200 µM
